# PD-1-Targeted IL-2v Expands a Novel Stem-like T Cell Subset Distinct from Anti-PD-1 Therapy to Enhance Effector Differentiation

**DOI:** 10.64898/2026.08.17.745170

**Authors:** Elzbieta Gralinska, Caterina Scirgolea, Amrita Manchala, Maria Karagianni, Greta Durini, Tamara Hüsser, Emilio Yangüez, Valeria Nicolini, Said Aktas, Laura Codarri Deak

**Author notes:** These authors contributed equally to this work (Said Aktas, Laura Codarri Deak).

## Abstract

Tumor-draining lymph nodes serve as critical sites for the generation, maintenance and differentiation of stem-like CD8⁺ T cells during antitumor immune responses. Recent evidence has shown that delivering interleukin-2 to PD1⁺ stem-like CD8⁺ T cells using PD1-IL2v, an immunocytokine combining PD-1 blockade and IL-2R agonism, promotes their differentiation into potent effector cells with enhanced tumor-killing capacity. However, it remains unclear how targeted interleukin-2 therapies imprint early stem-like T-cell differentiation programs. Here, using single-cell transcriptomics and T-cell receptor sequencing in murine pancreatic tumor models, we demonstrate that PD1-IL2v induces an early bifurcation in the differentiation of stem-like CD8⁺ T cells within tumor-draining lymph nodes. We identify an effector-primed stem-like population characterized by the expression of interferon-response genes, natural killer cell receptor genes, and Cx3cr1, consistent with activation of interleukin-2 and STAT5-associated programs. Clonal tracking revealed substantial overlap between these lymph node-derived cells and intratumoral effector populations, supporting a developmental relationship between early priming in lymph nodes and downstream effector differentiation. In contrast, an alternative stem-like state that displayed features associated with T-cell exhaustion, including increased Tox expression, was observed upon PD-1 therapy. Together, these findings identify an early branch point in stem-like T-cell differentiation and provide mechanistic insight into how PD1-IL2v circumvents exhaustion pathways to preferentially generate functional antitumor immunity.

## INTRODUCTION

Immune checkpoint inhibitors (ICIs) targeting the PD-1 and CTLA-4 pathways have marked a significant advancement in cancer treatment *(1–4)*. However, their clinical efficacy remains limited, mainly due to primary or adaptive resistance mechanisms, including the expression of additional non-redundant immune checkpoints (*5–7*). Other immunotherapy approaches include the clinical use of Interleukin-2 (IL-2), the first cytokine approved for the treatment of metastatic melanoma and renal cell carcinoma (*8–11*). Despite improving tumor cell killing, its therapeutic utility is hampered by systemic toxicity associated with high dosing and frequent administration needed to overcome the poor pharmacokinetic profile required to target CD8+ tumor-infiltrating lymphocytes (TILs) (*12–13*).

To overcome these limitations, engineered IL-2 based therapies have been a major focus point in the development of next generation immunotherapies. These IL-2 variants, such as βγ-biased derivatives, are engineered to provide improved safety and efficacy profiles. By reducing binding to the IL-2R α subunit (CD25), these variants minimize the detrimental targeting of immunosuppressive regulatory T cells (Tregs) and pulmonary endothelial cells, the latter of which causes vascular leak syndrome (*14–18*). In addition, strategies such as fusing IL-2 to immune- or tumor stroma-targeting antibodies have been explored to enhance localized delivery, thereby improving the on-target, off-tumor safety profile (*19–20*).

PD1-IL2v, the first immune-cell targeted IL-2 variant, was designed to selectively deliver IL-2 receptor agonism in cis to PD-1+ CD8+ T cells, while simultaneously blocking this immune checkpoint receptor (*16–17*). Its IL-2R binding moiety (IL2v), is engineered to be devoid of binding ability to CD25, thus avoiding unwanted IL-2 targeting of Tregs, thereby enhancing its therapeutic potential. A previously published pre-clinical study has described PD1-IL2v’s ability to preferentially target PD-1+TCF-1+ stem-like CD8+ T cells and promote their differentiation into a functionally superior effector progeny (PD-1+ TCF-1-), known as “better effector” cells, defined as being Granzyme B+ TIM-3- (*16*). These findings highlight that while anti-PD-1 predominantly drives the accumulation of terminally exhausted CD8+ T cells, PD1-IL2v redirects this differentiation towards a functionally and phenotypically superior effector fate. Whilst this differentiation has been described to occur within the tumor microenvironment, it remains unclear whether this lineage commitment is initiated locally or established at earlier stages of the immune response within tumor draining lymph nodes (TdLNs), which serve as reservoirs of stem-like CD8+ T cells.

To address this, we performed single-cell RNA sequencing (scRNA-seq), TCR sequencing (TCR-seq), and fate-mapping approaches on tumors and TdLN samples from a murine pancreatic mouse tumor model, identifying in the latter two transcriptionally distinct stem-like CD8⁺ T-cell populations. We show that muPD1-IL2v treatment promotes the expansion of a previously undescribed stem-like subset with an effector-primed transcriptional program, distinct from populations observed under other treatments.

Through a TCR clonotype tracking approach across TdLN and tumor compartments, we demonstrate that this muPD1-IL2v-induced stem-like population is clonally linked to tumor-resident better effector T cells, whereas the second stem-like population is associated with exhausted states of tumor-specific T-cells. These findings reveal an early divergence in stem-like T-cell differentiation within TdLNs, preceding tumor infiltration. Together, our results provide mechanistic insight into how PD1-IL2v reshapes CD8⁺ T-cell differentiation and promotes the generation of functionally superior anti-tumor responses.

## RESULTS

### PD1-IL2v elicits a novel stem-like T cell subset in TdLNs of Panc02-H7-Fluc mouse tumor

We previously reported the effects of muPD1-IL2v on stem-like CD8^+^ T cells and their progeny in the tumors of immunocompetent syngeneic mice challenged subcutaneously with a pancreatic adenocarcinoma cell line (Panc02-H7-Fluc) (*16*). We described that muPD1-IL2v elicited the differentiation of stem-like T cells towards a novel and unique subset of CD8^+^ TILs with heightened effector and anti-tumor functions. Given that stem-like T cells are mainly present in the TdLNs, we decided to investigate the effects of muPD1-IL2v treatment in this compartment. For this purpose, mice were either left untreated or treated with muPD1-IL2v, anti-muPD-1 and stroma-targeted muFAP-IL2v as monotherapies, or with a combination of the latter two as a combination therapy (Fig. 1A). Since the TIL data from this study have been reported previously (*16*), they are not reported here. Instead, we provide a novel analysis of previously unpublished TdLNs from the same experiment and interpret these findings in the context of the published TIL data. Applying scRNA-seq to TdLNs-derived CD8^+^ T cells enabled us the identification of five main T-cell subsets: naive, recently-primed, proliferating, stem-like, and a small population of exhausted CD8^+^ T cells. These findings provide new insights into the early stages of stem-like CD8^+^ T cell differentiation (Fig. 1, B and C).

**Fig. 1.**
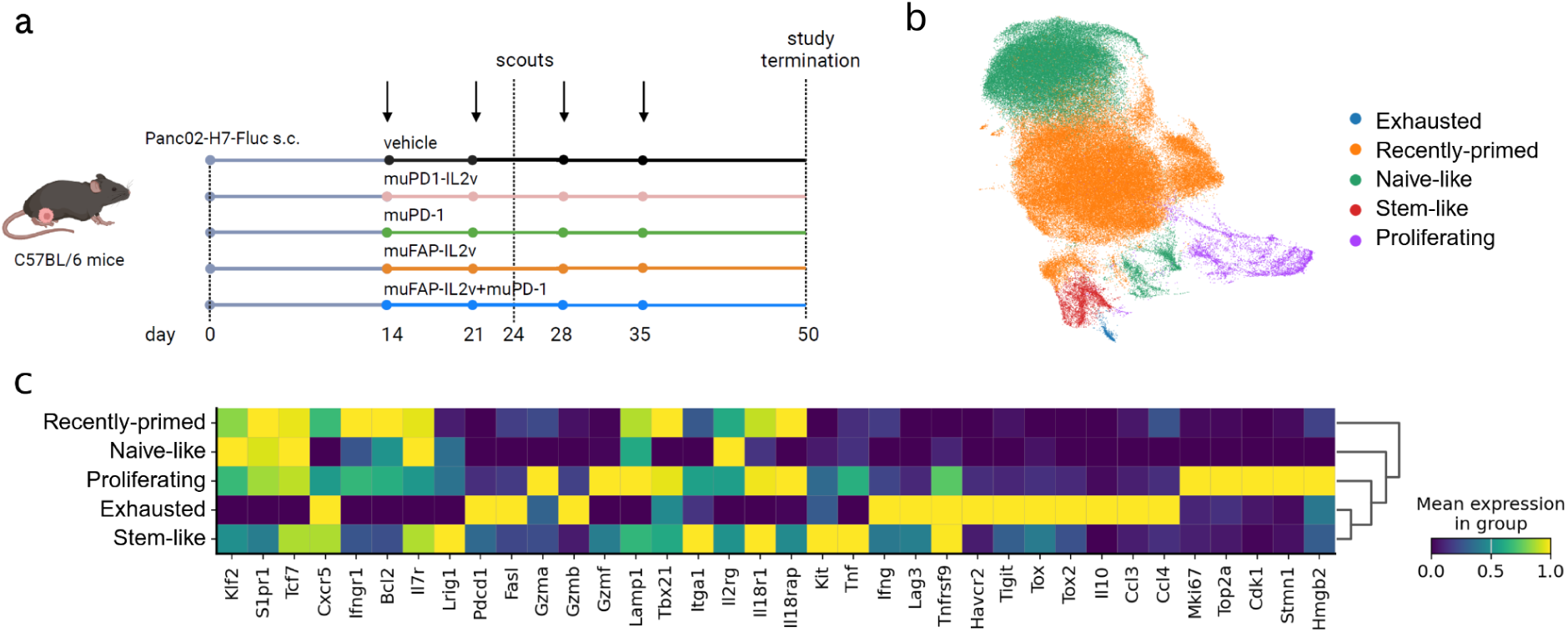
Analysis of single-cell RNA-seq data from CD8^+^ T cells from TdLNs from Panc02-H7-Fluc model. **(A)** Study design. **(B)** UMAP embedding with annotated CD8^+^ T cell types. **(C)** Expression of marker genes across annotated CD8^+^ T cell types.

Interestingly, we identified two distinct subsets of stem-like T cells in TdLN samples using single-cell fate mapping analysis of the scRNA-seq data. This analysis was performed using CellRank2 (*21–22*), a framework that constructs a cell-cell transition matrix based on transcriptional similarity, enabling the identification of both initial and terminal cell states, referred to as macrostates. CellRank2 further estimates cell fate probabilities, which quantify the likelihood of individual cells differentiating into specific terminal states. Using the inferred cell-cell transition probabilities, the framework can also simulate random walks that capture how cells may gradually progress through intermediate transcriptional states as they differentiate toward terminal states (Fig. 2A). Together, this approach provides insights into cellular state transitions within the biological system under investigation.

**Fig. 2.**
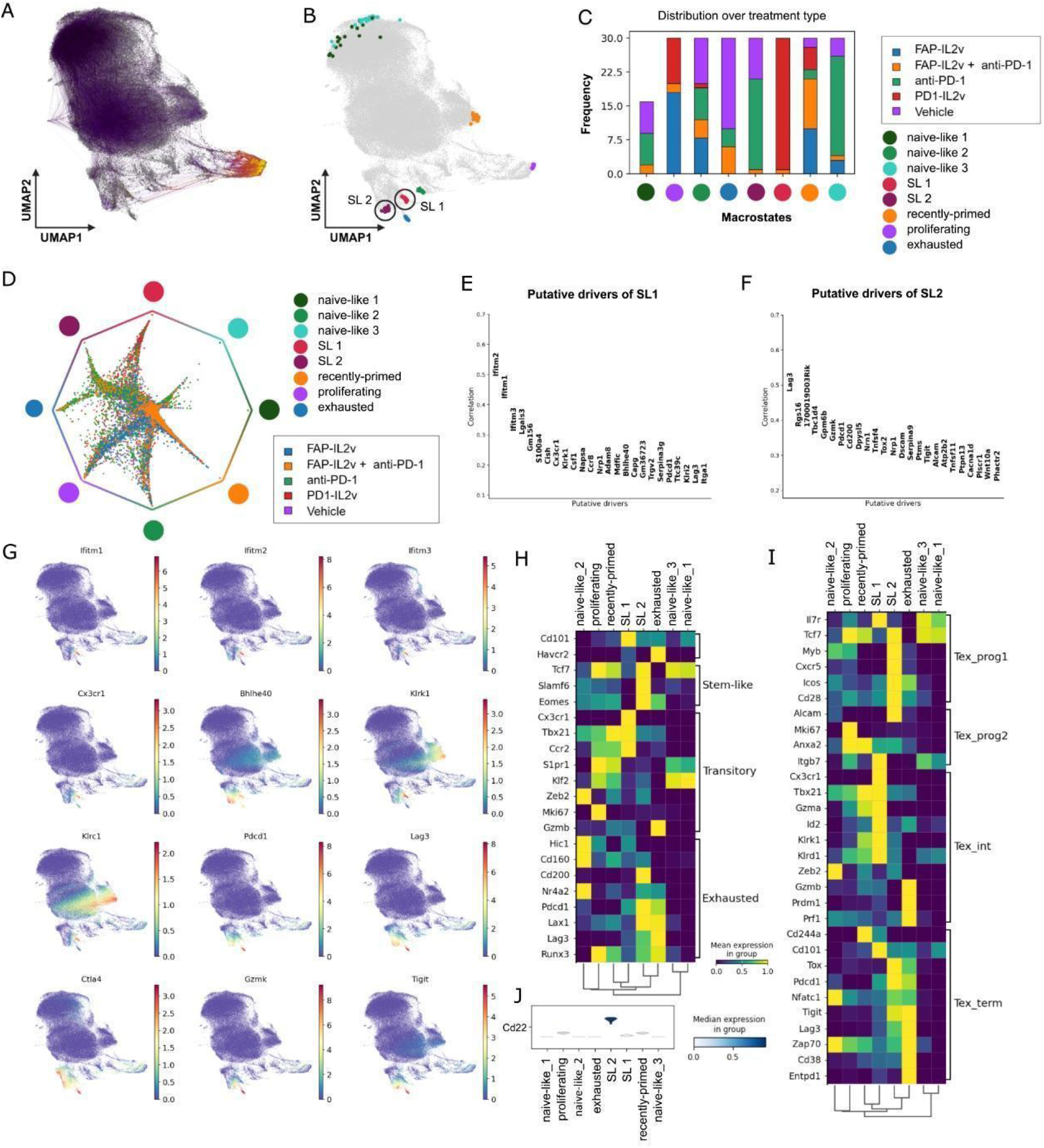
Single-cell fate mapping analysis of Panc02-H7-Fluc TdLN data. **(A)** UMAP embedding with random walks visualized. (**B)** UMAP embedding with eight inferred macrostates. Each color represents one macrostate. **(C)** Distribution of treatments across identified macrostates. **(D)** Circular projection representing cell fate probabilities toward each detected macrostate. The vertices of the polygon represent the detected macrostates. Each dot within the polygon represents one cell. Cells are colored by the treatment they originate from. The position of a given cell in the polygon represents its fate bias toward a given macrostate. (**E, F)** The putative driver genes detected for the (E) SL1 and (F) SL2 macrostates, respectively. Genes are ranked by the correlation between their expression value and fate probability toward a given state. **(G)** UMAP embeddings with genes characterizing SL1 and SL2 populations. **(H)** Expression of stem-like-, transitory- and exhausted-marker genes across SL1-, SL2-, and exhausted macrostates. **(I)** Expression of marker genes for Tex populations reported by Beltra et al. **(J)** Expression of Cd22 across the detected macrostates.

Two of the macrostates inferred by single-cell fate mapping analysis of the TdLN data were located within the stem-like T cell subset (Fig. 2B). Hereafter, we refer to these as “Stem-like 1” (SL1) and “Stem-like 2” (SL2) T cell populations. Analysis of the macrostate composition across treatment groups revealed a distinct pattern of cell distribution (Fig. 2C). Notably, the SL1 population was composed almost exclusively of cells from the muPD1-IL2v-treated mice and was absent in vehicle mice. This suggests that SL1 represents a treatment-induced differentiation rather than the expansion of a pre-existing population. Conversely, the SL2 population consisted predominantly of cells originating from anti-PD-1 treated and vehicle-treated mice. These findings suggest that muPD1-IL2v and anti-muPD-1 give rise to already transcriptionally distinct subsets of stem-like CD8^+^ T cells within TdLNs.

To further assess lineage priming, we visualized fate bias using a circular projection of cell fate probabilities (Fig. 2D). In this representation, terminal states are placed evenly around a unit circle, and individual cells are embedded within the circle according to their relative probability of transitioning toward each terminal fate. Consequently, the position of each cell reflects its putative lineage priming: the closer a given cell is to a terminal state, the greater the probability that it will transition toward that state. In this projection, the SL2 population is positioned closer to the exhausted T cell state than to SL1, indicating a stronger bias toward the exhausted lineage. Many cells lie along the axis between SL2 and the exhausted state, reflecting comparable transition probabilities and indicating closely related lineage priming. In contrast, SL1 cells are positioned further from this trajectory, consistent with weaker link to exhaustion.

### PD1-IL2v induces a transitory Cx3cr1+ subset within TdLNs

To gain further insight into the differences in transcriptional profiles between the SL1 and SL2 populations, we examined genes that may be driving the differentiation of these two macrostates (Fig. 2, E, F and G). The SL1 population exhibited a transcriptional profile enriched for genes associated with effector functions, including interferon response genes (*Ifitm1, Ifitm2, Ifitm3, Cish*) and NK-related genes (*Klrk1, Klri1, Klrc1*). Furthermore, SL1 cells showed increased expression of *Cx3cr1*, a marker of tumor-specific effector-memory T cells, and *Bhlhe40*, a transcription factor known to regulate anti-tumor T-cell effector responses. In contrast, the SL2 population was predominantly characterized by exhaustion-associated genes, including immune checkpoint receptors (*Lag3, Pdcd1, Ctla4*, and *Tigit*). Comprehensive lists of detected driver genes for the SL1 and SL2 macrostates, as well as other macrostates, are provided in Table S1, and a list of differentially expressed genes identified using the Wilcoxon test is provided in Table S2.

To further characterize the biological processes associated with these transcriptional differences, we performed Gene Ontology (GO) term analysis. We found an enrichment of distinct biological processes between the SL1 and SL2 populations. The SL1 population was enriched for pro-inflammatory and co-stimulatory processes, including: T-cell costimulation, inflammatory response, cytokine activity, C-X-C chemokine binding. In contrast, the SL2 population showed enrichment for immunoregulatory processes, such as: negative regulation of T cell migration, positive regulation of macrophage cytokines, and positive regulation of interleukin-10 production. Collectively, these results highlight a more pro-inflammatory transcriptional profile in SL1 cells, whereas SL2 cells express genes associated with anti-inflammatory and immunosuppressive programs.

The increased expression of *Cx3cr1* in the SL1 population raises the possibility that these cells are related to the transitory CD101⁻Tim3⁺ CD8⁺ T cell population reported by Hudson et al. (*23*) and the analogous CX3CR1⁺ population described by Zander et al. (2019) (*24*). To directly compare these populations, we evaluated the expression of gene signatures associated with stem-like, transitory, and exhausted states defined by Hudson et al., together with markers distinguishing Slamf6⁺ (Ly108⁺) (analogous to stem-like), CX3CR1⁺ (analogous to transitory), and Slamf6⁻CX3CR1⁻ (analogous to exhausted) subsets identified by Zander et al. (Fig. 2H).

Despite differences in experimental design between these studies, we observed the convergence of several transcriptional features. The SL1 population showed elevated expression of *Cx3cr1*, *Tbx21*, and *Ccr2*, supporting its similarity to the transitory state. However, *Cd101* expression was unexpectedly higher in SL1 than in the exhausted population, suggesting that SL1 more closely resembles the CX3CR1^+^ population reported by Zander et al., while differing from the CD101⁻Tim3^+^ subset described by Hudson et al. In contrast, the SL2 population exhibited a transcriptional signature more closely aligned with stem-like CD101⁻Tim3⁻ and Slamf6^+^ cells, suggesting an earlier differentiation state than SL1.

To further position these populations within established developmental frameworks of exhausted CD8^+^ T (Tex) cells, we compared our data with the transcriptional profiles defined by Beltra et al. (*25*), which distinguish two TCF1^+^ progenitor subsets (Tex^prog1^ and Tex^prog2^), an intermediate subset (Tex^int^), and a terminally exhausted subset (Tex^term^) (Fig. 2I). Analysis of the reported gene signatures across detected macrostates revealed that the SL1 macrostate most closely resembled the Tex^int^ population, including elevated expression of *Cx3cr1* and *Tbx21*. The SL2 macrostate showed similarity to the Tex^prog1^ and Tex^prog2^ populations, with partial expression of Tex^term^ signature genes (Fig. 2I). Additionally, we observed overexpression of *Cd22* in the SL2 population (Fig. 2J). Even though these comparisons suggest that, within the Tex differentiation framework, SL1 represents a different stage than SL2, it remains unclear whether SL1 and SL2 represent sequential stages of differentiation or distinct alternative populations. Despite the transcriptional similarities of the SL1 population to transitory and Tex^int^ cells, we detected SL1 cells in TdLNs, indicating that intermediate effector-like Tex states can arise or persist within lymphoid tissues.

### IL2-STAT5 axis may underlie the early divergence between SL1 and SL2

Recent work by Beltra et al. reported that, within the developmental hierarchy comprising Tex^prog1^, Tex^prog2^, Tex^int^, and Tex^term^ states (*25*), STAT5 signaling directs the transition from Tex^prog^to Tex^int^ cells and promotes the acquisition of an effector- and NK-like transcriptional program, characterized by genes such as *Cx3cr1, Tbx21, Gzma, Gzmb,* and *Klr* family receptors (*26*). In the same study, constitutive STAT5 activity (STAT5CA) was shown to antagonize *Tox* expression and reprogram exhausted CD8^+^ T cells toward a durable effector-like state, whereas loss of STAT5 impaired Tex^int^ differentiation and maintained cells in progenitor-like states. Beltra et al. further demonstrated that STAT5 signaling contributes to epigenetic remodeling during Tex differentiation, reshaping the chromatin landscape of exhausted CD8^+^ T cells toward effector- and memory-associated programs. Consistent with these observations, enhanced STAT5 activity increased the abundance of Tex^int^ cells and amplified responses to PD-L1 blockade, while re-engagement of STAT5 activity in Tex^prog^ cells partially reversed exhaustion-associated programs and restored effector functions. Collectively, these findings established the IL-2-Stat5 axis as a central regulator of intermediate Tex differentiation and epigenetic reprogramming.

Consistent with these reported findings, we observed that the SL1 population in the TdLNs exhibited high expression of *Il2ra* (CD25) (Fig. 3A), in line with its enrichment in PD1-IL2v-treated samples, which is expected during sustained IL-2R signaling to PD-1⁺ stem-like CD8⁺ T cells upon PD1-IL2v treatment. Although *Stat5a* expression was not increased at the transcriptional level in SL1 compared to SL2, this is expected given that STAT5 activity is primarily regulated through phosphorylation rather than mRNA abundance (*27*). Nevertheless, the transcriptional profile of SL1 cells was consistent with activation of STAT5-dependent programs, including elevated expression of *Cx3cr1, Tbx21*, and NK receptor genes (Fig. 3B). In contrast, the SL2 population showed increased levels of *Tox*, consistent with a more exhaustion-prone state. Given that both SL1- and SL2-like populations were identified within tumor-draining lymph nodes, this indicates that transcriptional programs corresponding to intermediate and progenitor Tex states are already present within lymphoid tissues. Together, these observations suggest that IL-2R signaling may underlie the difference between SL1 and SL2 states, with SL1 representing an IL-2-responsive, effector-primed population and SL2 retaining features of a progenitor-like, exhaustion-associated state.

**Fig. 3.**
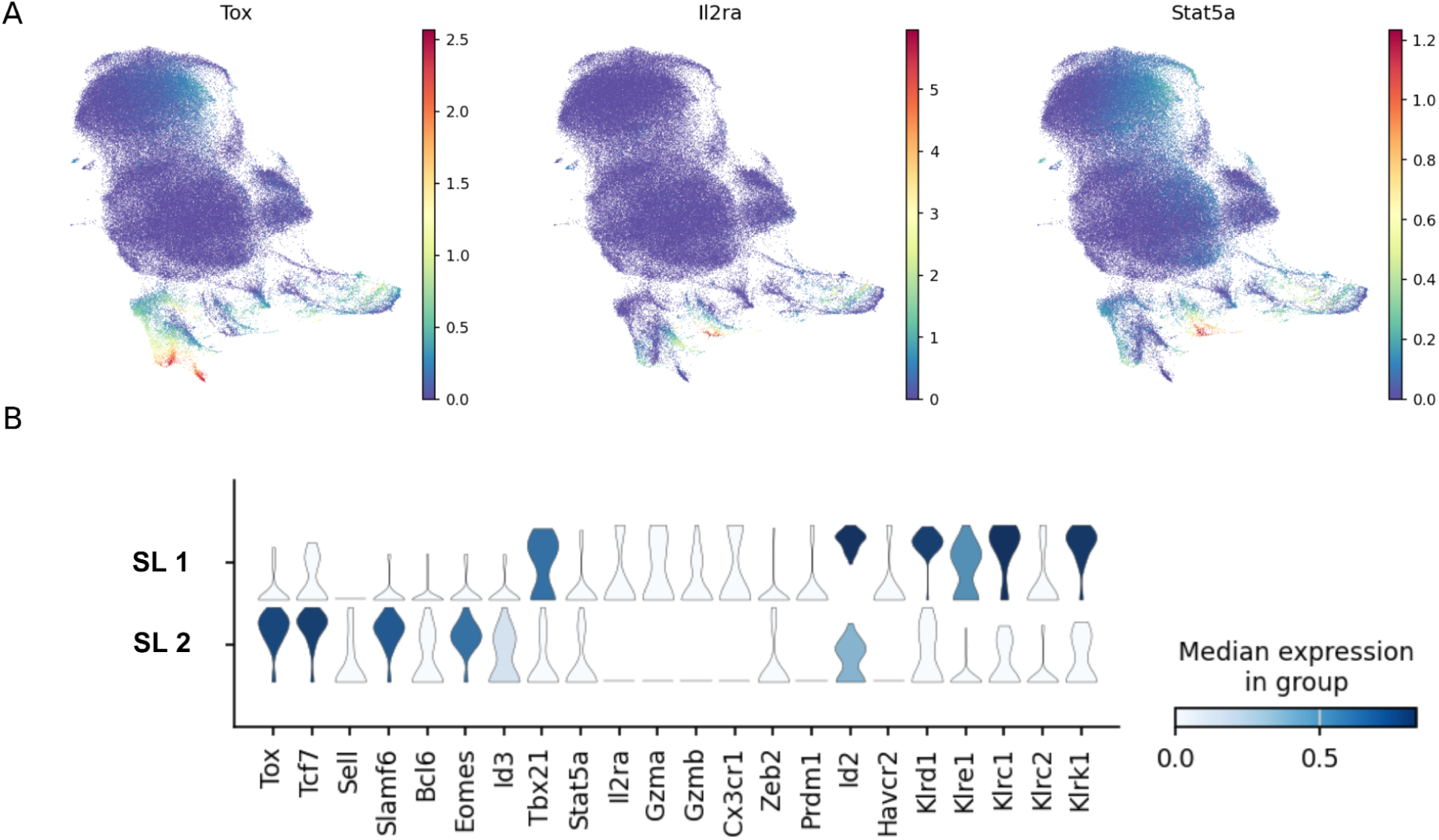
IL2-STAT5 axis-related genes in Panc02-H7-Fluc TdLN data. **(A)** UMAP embeddings representing expression of Tox, Il2ra (CD25), and Stat5a in CD8⁺ T cells TdLNs. (**B)** Violin plots representing the expression of signature genes in SL1 and SL2 populations.

### SL1 cells from TdLNs share their TCR repertoire with better effector cells in the tumor

TdLNs have been reported to act as reservoirs of stem-like T cells and represent the site of initial tumor-specific CD8⁺ T-cell differentiation prior to migration to the tumor site (*28–30*). Stem-like T cells generated in TdLNs subsequently traffic to the tumor site, where they can further differentiate.

Accordingly, our initial analyses focused on TdLN samples to investigate mechanisms governing early T cell differentiation. Our previous work demonstrated that systemic administration of muPD1-IL2v promotes the differentiation of stem-like T cells into “better effector” CD8^+^ T cells within the tumor, and that accumulation of these cells provided better therapeutic efficacy (*16*). Building on these findings, we investigated whether the SL1 or SL2 populations identified in TdLNs represent the primary precursor pool to the observed intra-tumoral better effector T cell populations. To address this question, we utilized paired TCR sequencing (TCR-seq) and scRNA-seq data from both TdLN and tumor samples, enabling clonotype tracking between these two sites. Clonotype analysis was performed based on CDR3 sequences and and the detected SL1- and SL2-associated clonotypes were projected onto the UMAP representation of CD8^+^ T cells from tumor to examine whether TdLN SL1 and SL2 populations are clonally related to distinct tumor-infiltrating T-cell populations.

Clonotype analysis revealed substantial TCR sharing between SL1 cells from TdLNs and intra-tumoral better effector T cells characterized by enhanced cytotoxic signatures and uniquely expanded by PD1-IL2v treatment (Fig. 4, A and B). Within this cluster, 8.80% of cells shared clonotypes with SL1 cells, whereas 1.23% shared clonotypes with SL2 cells, indicating stronger clonal overlap between better effectors in the tumors with the SL1 population from the TdLNs. In contrast, TCR sharing between SL2 cells and tumor-infiltrating populations was most pronounced in exhausted and memory/effector clusters. Specifically, 8.99% of exhausted cells and 3.19% of memory/effector cells shared clonotypes with SL2 cells, compared with 0.64% and 1.83%, respectively, for SL1 cells (Fig. 4C).

**Fig. 4.**
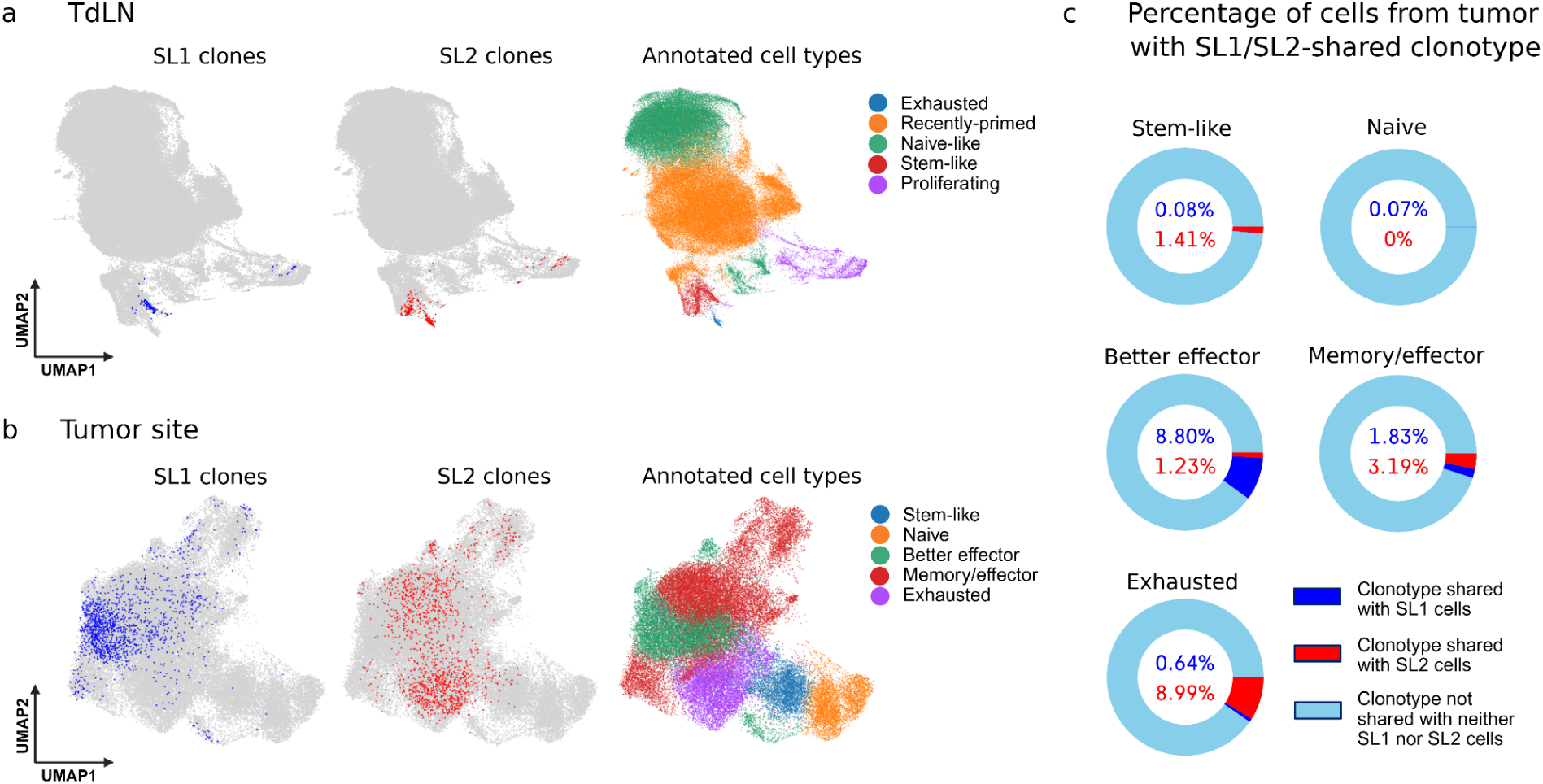
SL1- and SL2-derived clonotypes within TdLN and tumors. **(A, B)** UMAP embeddings representing cells sharing their clonotype with SL1- (blue) and SL2-derived (red) clonotypes in (A) dLN and (B) tumor data. The UMAP embeddings on the right side represent annotated cell types in both tissues. **(C)** Percentage of cells in tumor sharing their clonotype with TdLN SL1 and SL2 cells. Each pie chart represents one cell cluster from tumor. Numbers in the circles represent the percentage of cells from a given cell cluster, sharing their clonotype with TdLN SL1 (blue) and SL2 (red) cells.

Taken together, these patterns of clonotype overlap indicate distinct clonal relationships between TdLN SL1 and SL2 populations and specific intra-tumoral T cell clusters indicative of divergent differentiation pathways. These data support a model in which the TdLN SL1 population serves as a reservoir of highly functional better effector cells, whereas the SL2 population preferentially leads to the accumulation of terminally exhausted T cells within the tumor.

### Validation of the findings in the KPC-ADGPK tumor model

To validate the effects of muPD1-IL2v treatment on stem-like CD8⁺ T cells in TdLNs observed in the Panc02-H7-Fluc model, we performed an independent in vivo efficacy study using syngeneic C57BL/6 mice subcutaneously challenged with KPC 4662-ADPGK, a pancreatic ductal adenocarcinoma cell line expressing the ADPGK tumor antigen, used as a more physiological model to study the immune response towards an immunodominant tumor neoantigen. Analogous to the Panc02-H7-Fluc experiment, mice were treated with either PD1-IL2v and anti-PD-1, as monotherapies, or anti-PD-1 in combination with FAP-IL2v; vehicle-treated mice served as controls (Fig. 5, A-E).

**Fig. 5.**
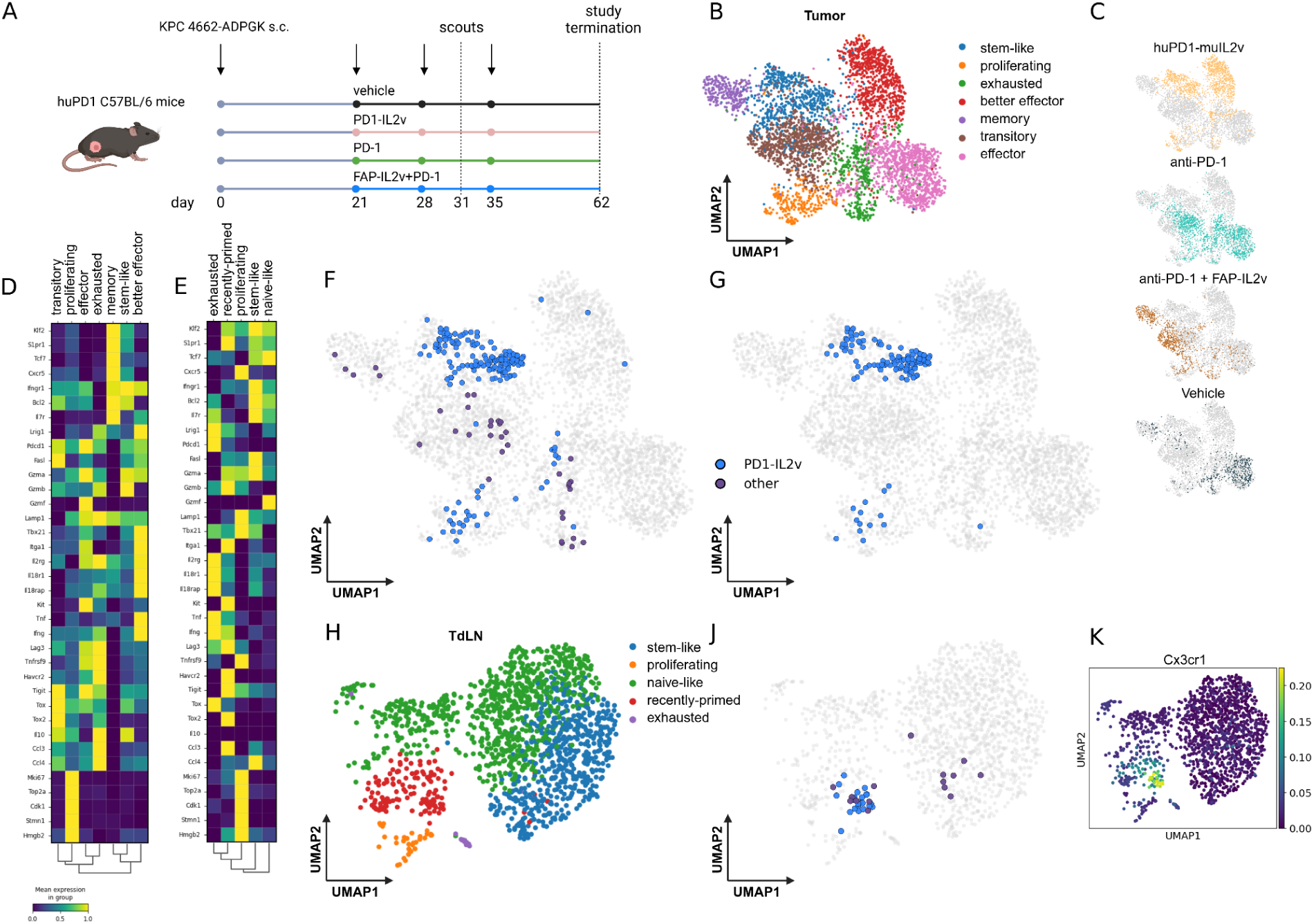
Clonotype analysis in the KPC ADPGK tumor model. **(A)** Study design. **(B)** UMAP embedding representing cell types in Tumor. **(C)** Distribution of treatments. Each UMAP embedding represents one treatment. **(D, E)** Matrixplots for (D) Tumor and (E) TdLN data. **(F)** UMAP embedding representing T cells in the tumor sharing their clonotype with TdLN cells. Clonotypes originating from PD1-IL2v-treated samples are highlighted in blue. Clonotypes from anti-PD-1 and anti-PD-1 with FAP-IL2v (called “other”) are in purple (blue). **(G)** UMAP embedding representing three most expanded clonotypes from (F). **(H)** UMAP embedding representing cell types in TdLNs. **(J)** UMAP embedding representing cells from TdLNs with the same clonotype as cells highlighted in (F). **(K)** UMAP embedding representing the expression of Cx3cr1 in TdLNs.

Paired scRNA-seq and TCR-seq data from tumor and TdLN samples enabled tracking of shared clonotypes across anatomical sites. This clonotype analysis of paired TdLN and tumor samples revealed two distinct groups of shared clonotypes. The first group comprised clonotypes originating from cells in the TdLNs upon PD1-IL2v-therapy (Fig. 5F). Within this group we found three clonotypes that were also strongly expanded in the tumor (Fig. 5G). The majority of TILs which shared these clonotypes belonged to the stem-like CD8^+^ T cell cluster, with only a minor fraction detected in proliferating and exhausted populations. Additional clonotypes present in PD1-IL2v-treated TdLN cells were less expanded and were distributed across stem-like, proliferating, better effector, and exhausted clusters, although at low frequencies. The second group consisted of cells derived from TdLNs upon either anti-PD-1 monotherapy or in combination with FAP-IL2v. This group showed limited expansion in the tumor and was primarily associated with transitory-, memory- and exhausted T cell clusters (Fig. 5, F, H and J).

To further define the cellular origin of these clonotypes, we examined their distribution within TdLN subsets. Clonotypes associated with the tumor-resident stem-like population predominantly originated from PD1-IL2v-treated TdLN cells and were almost exclusively derived from the CX3CR1⁺ CD8⁺ T-cell population. In contrast, clonotypes enriched in exhausted tumor T cells were more frequently found in mice receiving the other treatments (Fig. 5, C and D). Importantly, the CX3CR1⁺ TdLN population corresponds to the SL1 subset, suggesting that PD1-IL2v-responsive, CX3CR1⁺ CD8^+^ T cells in TdLNs may represent a major source of tumor-infiltrating clonotypes, particularly those contributing to the stem-like compartment within the tumor.

In conclusion, for the PD1-IL2v treatment we observed that, in contrast to the Panc02-H7-Fluc model, where TdLN-derived clonotypes were preferentially associated with TILs with the functional phenotype of better effectors, in the KPC-ADPGK model shared clonotypes in TdLN were predominantly linked to stem-like T cell clusters in the tumor, indicating that PD1-IL2v-driven differentiation trajectories depend on the intrinsic immunogenicity of the tumor antigens which vary across tumor models.

## DISCUSSION

The presence of stem-like CD8^+^ T cells has emerged as a key feature of adaptive immune response in chronic infection and cancer (*23, 31–33*). These cells function as a self-renewing reservoir capable of generating effector progeny in response to antigenic stimulation or immunotherapy. A detailed understanding of how immunotherapies modulate the early differentiation trajectories of these cells may therefore have important implications for improving treatment efficacy and refining patient enrichment strategies.

In this study, we have identified within TdLNs an early divergence of stem-like CD8^+^ T cells’ differentiation following PD1-IL2v treatment in the Panc02-H7-Fluc mouse tumor model. While our previous study has shown that cis-binding of PD1-IL2v to PD-1 and IL-2Rβγ promotes differentiation of stem-like T cells into better effector cells within the TME (*16*), our data extend this paradigm by demonstrating that these effects are already established at the level of TdLNs. Thus, PD1-IL2v acts by shaping early differentiation programs in lymphoid tissues.

In addition, our analyses further indicate that PD1-IL2v induces a distinct differentiation program in stem-like T cells, characterized by transcriptional signatures enriched for effector-associated genes, including interferon-response genes and NK receptor genes. This early transcriptional reprogramming within TdLNs is consistent with the better effector state observed in tumor-infiltrating CD8^+^ T cells.

By integrating TCR sequencing with single-cell transcriptomics data, we demonstrate that stem-like CD8^+^ T cells arising in TdLNs following muPD1-IL2v treatment share a substantial portion of their TCR repertoire with better effector T cells in the tumor. This clonal overlap supports a developmental continuum in which PD1-IL2v-primed stem-like T cells migrate from TdLNs to the tumor and differentiate into better effectors. In contrast, stem-like T cells generated under anti-PD-1 treatment showed great clonotype overlap with exhausted TILs and minimal overlap with the better effector population, indicating that muPD1-IL2v uniquely promotes this differentiation program. These findings provide a mechanistic explanation for the expansion of better effector T cells in PD1-IL2v-treated tumors, consistent with the previous study (*16*).

Interestingly, we observed model-dependent differences in the clonal relationships between TdLN and intratumoral T-cell populations. In the Panc02-H7-Fluc model, TdLN-derived SL1 cells shared clonotypes with better effector cells in the tumor, consistent with a scenario in which IL-2-responsive stem-like T cells generated in the TdLN contribute to TILs effector differentiation. In contrast, in the KPC-ADPGK model, clonotype overlap was predominantly detected between PD1-IL2v-responsive cells and the intratumoral stem-like population, with comparatively limited overlap with the better effector population. These findings suggest that, in the KPC model, antitumor responses may rely more heavily on local differentiation within the TME than on the continuous recruitment of newly generated T cells from the TdLN, with PD1-IL2v acting directly on an existing intratumoral progenitor pool. This interpretation is consistent with a previous work showing that in KPC tumors therapeutic efficacy is maintained despite FTY720-mediated inhibition of T cell egress from lymph nodes (*34*), indicating that in this model continuous recruitment of TdLN-derived cells is not essential for tumor control. Together, these observations indicate that the relative contributions of TdLN-derived and intratumoral T-cell populations to the therapeutic response are context-dependent and vary across tumor models.

Mechanistically, our data point to a central role for the IL-2-STAT5 axis in shaping this early divergence in stem-like T-cell fate. We observed that the SL1 population exhibits high expression of *Il2ra* (CD25), consistent with CD25 upregulation as the positive feedback loop following IL-2 receptor signaling in response to PD1-IL2v. Although *Stat5a* expression was not increased at the transcriptional level, this is expected given that STAT5 activity is regulated primarily through phosphorylation rather than mRNA abundance (*27, 35–36*). Consistent with this, SL1 cells displayed transcriptional features indicative of STAT5-dependent programs, including elevated expression of *Cx3cr1*, *Tbx21*, and NK receptor genes. In contrast, the SL2 population showed higher expression of *Tox* and progenitor-associated markers such as *Slamf6*, consistent with a more exhaustion-prone state.

In addition, SL2 cells selectively express CD22, an inhibitory receptor recently reported on subsets of progenitor exhausted CD8⁺ T cells. Recent work has linked CD22 expression to maintenance of a restrained progenitor-like state with limited differentiation toward effector populations during chronic antigen stimulation (*37*). The enrichment of CD22 within SL2 therefore further supports the interpretation that this population represents a more quiescent and exhaustion-associated stem-like state relative to SL1. Conversely, the absence of CD22 expression in SL1 is consistent with the effector-associated transcriptional program induced by PD1-IL2v treatment.

These observations align with prior work demonstrating that STAT5 signaling promotes differentiation toward intermediate effector-like Tex states while antagonizing TOX-driven exhaustion programs (*26*). Together, our data support a model in which PD1-IL2v-mediated IL-2R signaling promotes effector differentiation of stem-like CD8⁺ T cells through STAT5 activation and suppression of exhaustion-associated programs. More broadly, IL-2 signaling is known to tune CD8⁺ T-cell fate in a dose- and context-dependent manner, with stronger IL-2R signaling promoting effector differentiation and weaker or transient signaling favoring memory or exhaustion. In this context, the targeted and sustained IL-2R engagement together with PD-1 blockade provided by PD1-IL2v may shift this balance toward an effector-primed trajectory, consistent with the emergence of the SL1 population. Thus, cis- IL-2R agonism together with PD-1 blockade may represent a key upstream mechanism driving the divergence between SL1 and SL2 states.

Collectively, our findings uncover an early checkpoint in stem-like CD8⁺ T-cell differentiation within TdLNs following PD1-IL2v treatment. This previously unrecognized stage of lineage divergence links IL-2 signaling to the generation of better effector progeny, providing new insight into the mechanisms underlying responses to combination therapies. Importantly, similar differentiation patterns were observed in an independent dataset from a related IL-2-based therapy, a PD1-targeted IL-2 variant analogous to PD1-IL2v (*38*). This molecule, designed to deliver IL-2R signaling to PD-1⁺ T cells through a distinct engineering approach, shares key functional features with PD1-IL2v. Analysis of scRNA-seq data from MC38 tumor models treated with this agent identified a T-cell population resembling the SL1 subset, characterized by elevated expression of *Cx3cr1*, *Bhlhe40*, NK-related genes such as *Klrk1* and *Klrc1*, and interferon response genes (*Ifitm1* and *Ifitm2*). These observations raise the possibility that the emergence of effector-primed, CX3CR1⁺ stem-like populations could represent a broader feature of PD-1-targeted IL-2 therapies. Although this data was not the primary focus of the present study, it suggests that the emergence of effector-primed, CX3CR1⁺ stem-like populations may not be restricted to a single therapeutic modality, but instead reflect a more general mechanism of targeting both pathways simultaneously on the same cell allow for the engagement of the IL-2-STAT5 axis to reprogram stem-like CD8⁺ T cells toward effector differentiation while limiting exhaustion.

Finally, while our findings provide important insights into early T-cell differentiation in murine tumor models, future studies will be required to determine whether similar early bifurcation events occur in human tumors. Such efforts may help explain variability in clinical responses to immunotherapy and inform the development of more precise patient enrichment strategies.

## MATERIALS AND METHODS

### Mice

#### Panc02-H7-Fluc

As previously published, six- to 8-week-old female C57BL/6J were purchased from the Jackson Laboratory. All animal experiments were performed in accordance with National Institutes of Health and Emory University Institutional Animal Care and Use Committee guidelines.

#### KPC ADPGK

Six- to 8-week-old female C57BL/6J were purchased from the Jackson Laboratory. Mice were hosted in individually ventilated cages. All animal experiments were performed in accordance with National Institutes of Health and Emory University Institutional Animal Care and Use Committee guidelines.

### Mouse tumour models

#### Panc02-H7-Fluc

To test the in vivo efficacy of muPD1-IL2v compared with pembrolizumab, a subcutaneous syngeneic model was used in human PD-1-transgenic C57BL/6J mice (University of Oxford). In brief, 6- to 8-week-old female C57BL/6J mice (Charles River) were inoculated with 1 × 10^5^ Panc02-H7-Fluc cells injected subcutaneously.

Mice were maintained under specific-pathogen-free conditions with daily cycles of 12 h light/12 h darkness according to guidelines (temperature of 22 °C, dark/light cycle of 12 h and humidity of 50%; GV-SOLAS, FELASA), and food and water were provided ad libitum. Continuous health monitoring was carried out, and the experimental study protocol was reviewed and approved by the Veterinary Department of Canton Zurich.

Mice were randomized into different treatment groups, and therapy started when tumors reached an average volume of 200 mm3 as measured by caliper. All treatments were administered intravenously, and the following doses were investigated: muPD1-IL2v at 0.5 mg kg^−^, muFAP-IL2v at 2.5 mg kg^−^, muPD1 at 10 mg kg^−^ twice daily.

#### KPC ADPGK

To test the in vivo efficacy of PD1-IL2v compared with pembrolizumab, a subcutaneous syngeneic mouse model was used in human PD-1-transgenic C57BL/6J mice (University of Oxford). In brief, 6- to 8-week-old female C57BL/6J mice (Charles River) were inoculated with 3 × 10^5^ KPC 4662 cells injected subcutaneously.

To test the in vivo efficacy of PD1-IL2v compared with pembrolizumab or the combination of FAP-IL2v plus pembrolizumab, a subcutaneous syngeneic mouse model was used in human PD-1-transgenic C57BL/6J mice (University of Oxford). In brief, 6- to 8-week-old female C57BL/6J mice (Charles River) were inoculated with 3 × 10^5^ KPC 4662-ADPGK cells injected subcutaneously.

Mice were maintained under specific-pathogen-free conditions with daily cycles of 12 h light/12 h darkness according to guidelines (temperature of 22 °C, dark/light cycle of 12 h and humidity of 50%; GV-SOLAS, FELASA), and food and water were provided ad libitum. Continuous health monitoring was carried out, and the experimental study protocol was reviewed and approved by the Veterinary Department of Canton Zurich.

Mice were randomized into different treatment groups, and therapy started when tumors reached an average volume of 200 mm^3^ as measured by caliper. All treatments were administered intravenously once per week (with the exception of FAP-IL2v that was administered intraperitoneally), and the following doses were investigated: murinized PD1-IL2v at 0.5 mg kg^−^, murinized pembrolizumab at 3 mg kg^−^ and murinized FAP-IL2v at 1 mg kg^−^.

### Lymphocyte isolation

#### Panc02-H7-Fluc

Mice were euthanized according to animal welfare guidelines; tumor tissue and draining lymph nodes were isolated in the animal facility. Tumor tissue was transferred to PBS and was disrupted using manual scissors and the Miltenyi Gentle MACS machine. Subsequently, it was digested for 30 minutes at 37 °C in an enzyme mix consisting of RPMI with 10 mg ml^−^ DNase (Sigma-Aldrich) and 0.25 mg ml^−^ Liberase (Sigma-Aldrich). After digestion, the tissue mix was filtered through a 70-µm filter and resuspended as a single-cell suspension with an appropriate volume for subsequent staining with fluorescently labelled antibodies. Lymphocytes were mechanically isolated from draining lymph nodes with a pestle, filtered through a 70-µm filter and resuspended as a single-cell suspension with an appropriate volume for subsequent staining with fluorescently labelled antibodies.

#### KPC ADPGK

Mice were euthanized according to animal welfare guidelines; tumor tissue, draining lymph nodes and blood were isolated in the animal facility. Tumor tissue was transferred to PBS and was disrupted using manual scissors and the Miltenyi Gentle MACS machine. Subsequently, it was digested for 30 minutes at 37 °C in an enzyme mix consisting of RPMI with 10 mg ml^−^ DNase (Sigma-Aldrich) and 0.25 mg ml^−^ Liberase (Sigma-Aldrich). After digestion, the tissue mix was filtered through a 70-µm filter and resuspended as a single-cell suspension with an appropriate volume for subsequent staining with fluorescently labelled antibodies. Blood was transferred to heparin tubes, and red blood cells were lysed with an erythrocyte lysis buffer. After red blood cell lysis, cells were resuspended as a single-cell suspension with an appropriate volume for subsequent staining with fluorescently labelled antibodies. Lymphocytes were mechanically isolated from draining lymph nodes with a pestle, filtered through a 70-µm filter and resuspended as a single-cell suspension with an appropriate volume for subsequent staining with fluorescently labelled antibodies.

### Cell sorting

#### Panc02-H7-Fluc

Tumors and draining lymph nodes were digested as previously described and 1–10 x10^6^ cells were stored in liquid nitrogen. After thawing a batch of samples, cell suspensions from 3–5 tumours of the same treatment group were stained with the antibodies listed in Table 1; cells were kept on ice during the staining and the sorting procedures. Discrimination of living cells from dead cells was performed using Live/Dead APC-Cy7 (eBioscience, 65-0865-14; 1:500) . Cells were washed twice, filtered through a 40-µm cell strainer and sorted on a FACSAria III instrument (to enrich viable single CD45^+^CD11c-CD4-CD8^+^T cells).

### KPC ADPGK

Tumors and draining lymph nodes were digested as previously described and 1–10 x10^6^ cells were stored in liquid nitrogen. After thawing a batch of samples, CD45^+^ cells positive enrichment was performed using CD45 (TIL) MicroBeads (Miltenyi) and enriched CD45^+^ cells were stained with the antibodies listed in Table 1; cells were kept on ice during the staining and the sorting procedures. Discrimination of living cells from dead cells was performed using LIVE/DEAD™ Fixable Aqua Dead Cell dye (Invitrogen, L34957 ; 1:1000). Cells were washed twice, filtered through a 40-µm cell strainer and sorted on a FACSAria III instrument (to enrich viable single CD45^+^CD11b^-^CD3^+^ T cells).

### RNA isolation and RNA-seq

#### Panc02-H7-Fluc

Samples were randomized and processed in four different batches with eight samples each (tumours and lymph nodes were processed separately). Cells from tumors and draining-lymphnodes (n=3-5/each treatment group) were labeled with fluorescently conjugated monoclonal antibodies, FACS-sorted, washed resuspended in 1 ml PBS−/− containing 0.04% BSA. The cell number and viability of the sorted cells was determined using a Nexcelom Cellometer Auto 2000 and a total of up to 10,000 viable cells per sample were loaded into the 10x Genomics Chromium Connect Instrument. cDNA and library preparation was performed according to the manufacturer’s indications (10x Genomics Chromium Next GEM Automated Single Cell 5’ Reagent Kits v2 with TCR and feature barcoding) and the resulting libraries were sequenced in an Illumina NovaSeq6000 sequencer according to 10x Genomics recommendations (R1 = 26, i7 = 10, i5 =10, R2 = 90) to a depth of approx 20,000 reads per cell for the GEX library and 5,000 reads per cell for both the TCR and feature barcoding libraries. The list of antibodies labelled with a unique oligonucleotide tag identifier for CITE-seq is provided in Table S3.

### KPC ADPGK

Samples were randomized and processed in four different batches with eight samples each. Isolated CD45^+^ cells from tumors and draining-lymphnodes (n=4/each treatment group) were labeled with fluorescently conjugated monoclonal antibodies, FACS-sorted, washed resuspended in 1 ml PBS−/− containing 0.04% BSA. The cell number and viability of the sorted cells was determined using a Nexcelom Cellometer Auto 2000 and a total of up to 10,000 viable cells per sample were loaded into the 10x Genomics Chromium Connect Instrument. cDNA and library preparation was performed according to the manufacturer’s indications (10x Genomics Chromium Next GEM Automated Single Cell 5’ Reagent Kits v2 with TCR and feature barcoding) and the resulting libraries were sequenced in an Illumina NovaSeq X Plus sequencer according to 10x Genomics recommendations (R1 = 26, i7 = 10, i5 =10, R2 = 90) to a depth of approx 20,000 reads per cell for the GEX library and 5,000 reads per cell for both the TCR and feature barcoding libraries. The list of antibodies labelled with a unique oligonucleotide tag identifier for CITE-seq is provided in Table S3.

### Single-cell RNA, protein and TCR sequencing analyses

Raw FASTQ files were processed with CellRanger (count and vdj, v60.0.0) and aligned to the mouse reference transcriptome (mm10-2020-A), using the parameter ‘--expect-cells = 6000’. Cells with more than 200 detected counts were retained and merged across all samples for downstream analysis in Python using Scanpy (*39*) version 1.9.8 standard workflow. For Panc02-H7-Fluc samples, quality-control filtering was applied using the following parameters: min_genes = 100, min_cells =3, n_genes_by_counts < 6000 and >1000, pct_counts_mt < 8. For KPC ADPGK TdLN samples, quality-control filtering was applied using the following parameters: min_genes = 1,490, min_cells =2, n_genes_by_counts < 23,637 and >1000, pct_counts_mt < 2,25. For KPC ADPGK tumor samples, quality-control filtering was applied using the following parameters: min_genes = 967, min_cells =2, n_genes_by_counts < 26,247 and >1000, pct_counts_mt < 1,83. RNA counts were normalized using sc.pp.normalize using default parameters, followed by selection of highly variable genes. Total transcript counts and mitochondrial read content were regressed out prior to dimensionality reduction by PCA The first 50 principal components were used to construct the nearest-neighbor graph, perform Leiden clustering, and generate UMAP embeddings for visualization. Cell-type annotation was initially carried out using the Besca sig-annot module. CD8⁺ T-cell subsets were then refined based on the expression of RNA and protein markers together with gene-signature enrichment scores calculated using scanpy.tl.score_genes. Subsequent analyses were restricted to clusters containing CD8⁺ T cells. The macrostate analysis was conducted using CellRank2 version 2.0.0. Wilcoxon results for each macrostate were computed using sc.tl.rank_genes_groups with default parameters.

TCR-seq data analysis was performed in Python with Scirpy (*40*), and clonotypes were defined based on CDR3 sequence identity using the parameters: receptor_arms = "all" and dual_ir = "any".

### Data and code availability

The scRNA-seq, CITE-seq, and TCR-seq datasets generated from the KPC-ADPGK model and from tumor-draining lymph node samples of the Panc02-H7-Fluc model will be available upon request. TCR-seq data from the corresponding tumor samples from the Panc02-H7-Fluc model are available in ArrayExpress under accession number E-MTAB-11773.

### List of Supplementary Materials

Tables S1 to S3

### Funding

This study was funded by F. Hoffmann-La Roche Ltd. As employees of the funder, the authors were responsible for the study design, data collection, analysis, interpretation, the writing of the manuscript, and the decision to submit the paper for publication. The funder did not influence the results/outcomes of the study despite author affiliations with the funder.

### Author contributions

Conceptualization: EG, LCD, SA

Methodology: EG, CS, AM, GD, MK, VN, SA, LCD

Investigation: EG, AM, CS, GD, MK, TH, EY

Visualization: EG, CS

Analysis: EG

Project administration: EG

Supervision: LCD, SA Writing – original draft: EG

Writing – review & editing: EG, AM, CS, LCD, SA

## Acknowledgements

The authors gratefully acknowledge their colleagues at Roche pRED for their valuable contributions to this work. In particular, we thank Brigita Urben for processing the KPC-ADPGK sequencing data, Dzhansu Hasanova for her assistance with data upload, and Jehad Charo for his insightful discussions and continued support throughout the project.

